# On phylogenetic branch lengths distribution and the late acquistion of mitochondria

**DOI:** 10.1101/064873

**Authors:** Alexandros A. Pittis, Toni Gabaldón

## Abstract

In a recent article, Martin et. al. [1] criticize several methodological aspects of our recent study on the timing of the acquisition of mitochondria [2]. Here we show that our results are independent of the model-based partitioning of the data. In addition, we assess the robustness of our inferences by analyzing data from Martin et. al.

---

We first want to note that none of the points raised by Martin et. al. [1] affect the core of our conclusions - *i.e*. that differences in stem lengths relate to phylogenetic origin of LECA families so that they are shorter in bacterial, and particularly alpha-proteobacterial derived families - because the observed relationships i) are independent of the clustering performed in Figure 1 of [2], and ii) their criticism focuses on one single comparison of a single dataset but the differences are present across several datasets and approaches, including the very same dataset from the authors mentioned in their letter [3], as we show below. Secondly, their interpretation of our stem length measurement and how they extrapolate to branches subtending eukaryotic clades is conceptually flawed, as we also demonstrate below. Thus none of their arguments compromise at any rate the main conclusions of our article. We nevertheless want to discuss their points.

**Figure I Ascomycota stem length analysis.**
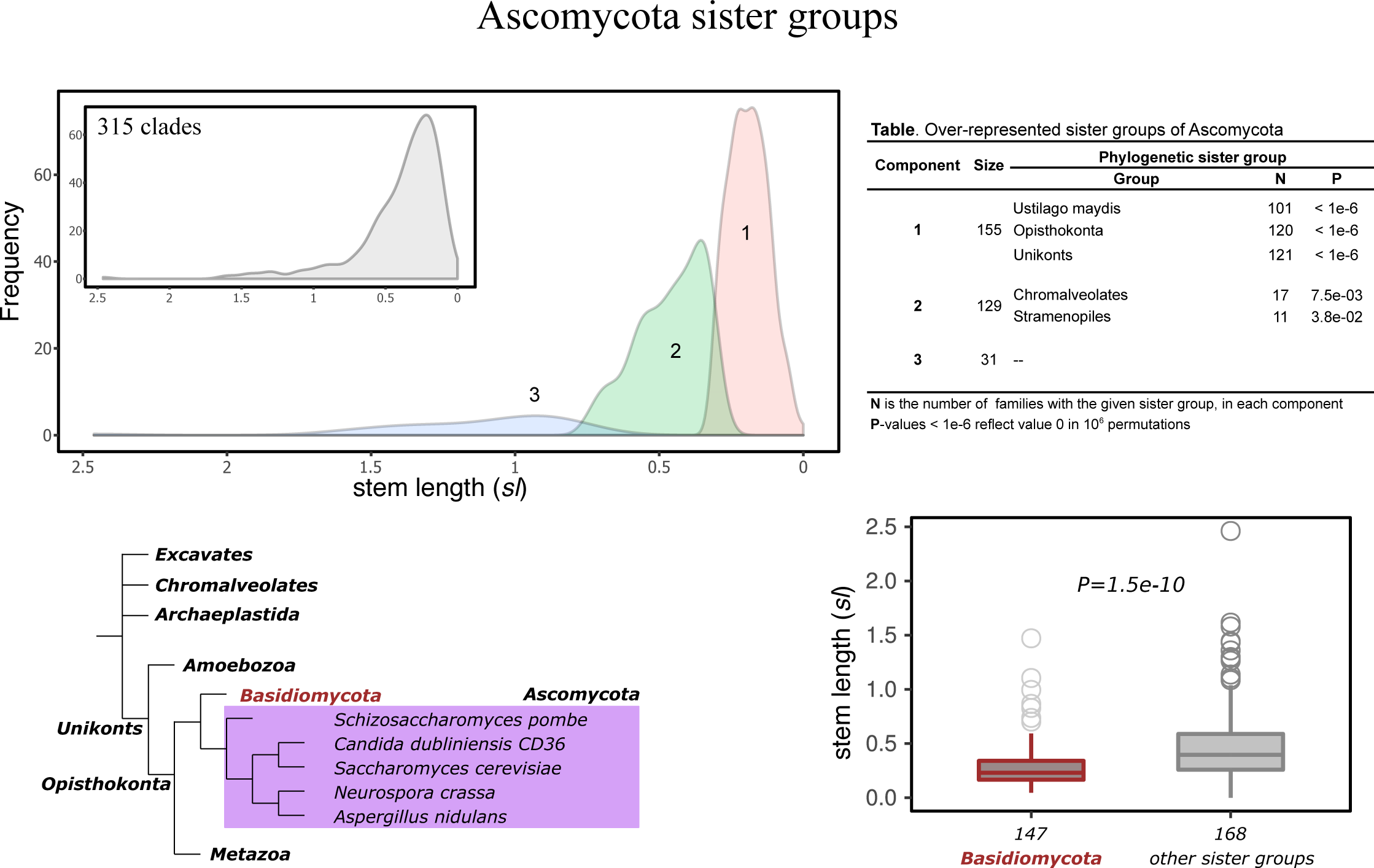
Different phylogenetic sister groups show significant differences in stem lengths according to their divergence times from Ascomycota. Gene losses in the sister group lineage can explain the alternative tree topologies and differences in estimated stem lengths.

Contrary to what Martin et al. [1] claim we do not assume a normal distribution of the global distribution of stem lengths. The claim that our statistical analyses are inappropriate is simply not true, we clearly explain all the methods used, and the tests performed to support observed differences are all nonparametric, without any assumption of normality. In Figure 1 we did use a probabilistic clustering method that fits a Gaussian mixture model, a mixture of normal distributions, assuming multimodality in the data. Martin et. al. [1] show that a unimodal log-normal distribution would better fit the data when the number of parameters is penalized. Does this demonstrates that the underlying distribution is not a composite of five gaussians? No, because when data are drawn from a five gaussian distributions with the obtained parameters, in 81% of the cases a log-normal distribution would be (wrongly) preferred using the BIC criterion. Also, the fact that any randomly sampled log-normal distribution could be fitted by a mixture model is by no means a surprise. In fact any distribution of data could be fitted by a finite number of mixture components, and this is precisely why these mixture models are commonly used as universal function approximators and as a tool to partition various kinds of data. Finally the definition of overfitting is not BIC inflation but the lack of predictive power. Thus other parameters have to be considered when assessing whether a model provides a reasonable representation of the data. The use of the EM algorithm is justified as a method for partitioning the data because i) we may expect composite of signals from a proteome (LECA) with at least two ancestral components (Archaeal host, and bacterial endosymbiont), and ii) prior studies have suggested that normalized branch lengths measurements as the ones used here to be approximately normal [4]. The assumtion of a unimodal distribution such as the one proposed by Martin et. al. [1] does not capture the expected mixture origins for a chimeric proteome and does not fit with the observation that differences in stem lengths relate to non-homogeneous phylogenetic origins. In any case our results are independent of this clustering exercise as the differences in stem lengths are apparent when simply grouping the LECA families according to their sister clades (Fig. 2 and Extended Data Fig. 1b of [2] or when using other forms of clustering the data such as equal binning (results not shown).

Their purported extrapolation of our analyses to eukaryotic clades and their derived dates is totally flawed and misleading. First of all, we explicitly say that we do not assume constant rates *(i.e*. molecular clock), and our normalized branch length is a measurement that is proportional to time but multiplied by a ratio between the rate preceding and postdating LECA, so their timing exercise, providing date estimates, is completely ungrounded. Secondly, Martin et al. [1] consider the normalized *sl* to yield arbitrary values, resulting in a log-normal distribution. This openly contradicts the observation that families of different prokaryotic origins show significant differences in *sl* and also *rsl* values. All our analyses robustly prove the opposite, there are differences and these differences reflect the relative divergence times. The cases of the cyanobacterial signal in Archaeplastida (Extended Data Fig. 3 of [2]) and of Lokiarchaeota signal in LECA (Extended Data Fig. 7 of [2]) nicely indicate the validity of the measurement. Expecting some extreme *ebl* values to reflect radical adaptations and fast rates of some lineages, we used the median because of its robustness with respect to extreme outliers (see Methods of [2]). We also tried not accounting for fast evolving taxonomic groups in the calculations, without any change in our main results. All these observations are not explained by the interpretation of the data provided by Martin. et. al. [1]. Furthermore, Martin et. al. [1] show that the normalized branch lengths sub-tending each eukaryotic clade follow log-normal distributions, and conclude that this observation demonstrates that this is natural variation for branches meant to represent a single time interval (e.g. divergence of fungi from metazoans). By adopting this assumption they are surprisingly ignoring that eukaryotic families are also subject to differential gene loss and other processes, which would result in multiple underlying patterns of the sub-tending branches (i.e. the sub-tending branch of a fungal family, which was lost in metazoans does not derive from the divergence between fungi and metazoans, but from the deeper divergence of fungi and other unikonts). This becomes apparent when controlling for the relationship of the normalized branch lengths with the phylogenetic affiliation of the sister branch - a key step in our analyses which they ignore. Indeed applying to the eukaryotic clades an EM-based clustering and measuring enrichments in phylogenetic affiliations as we did in our previous analysis [2] reveals major underlying distributions related with the nature of the sister group (Fig. 1). Thus, in this case also, the variation of *sl* values, interpreted by the authors as “vividly documenting abundant branch length variation”, is clearly shown to naturally carry the signal of different divergence times. So yes, the *sl* values in eukaryotic groups do imply phases of early and late divergence times due to gene loss or other biological events, as they do in the case of LECA. Of note this is a new, independent demonstration that variation in stem lengths relate with underlying variation in phylogenetic distribution, and provides additional support to our approach.

Finally, Martin et. al. [1] Focus their criticism in only one of our comparisons and on only one of the datasets used. For that dataset, they wrongly claim that we reused eukaryotic sequences in the different tree. This is false. Given the multidomain nature of eukaryotic protein sequences, the source of that dataset [5] may incorporate a given protein to more than one orthologous cluster. However we made sure we only used the orthologous sequence regions in a given analysis, thus never re-using a given eukaryotic sequence. Our analyses use standard filtering approaches but they claim that statistical significance for one of our comparisons (alpha-proteobacterial to other bacteria) is lost when applying additional ad hoc filtering on top of our previous filtering steps. We must note that even applying their filterings and using a permutation test as the one used in our paper, the alpha-proteobacterial *sl* values, remain significantly lower compared to other bacteria (P=1e−2, accounting only for families with eukaryotic sequence lengths >= 100 and P=3.7e−2, accounting only for alignments with gaps <= 50%, 10^6^ permutations). The loss of significance in some of the tests when artificially reducing the data is unsurprising. We are focusing on very ancient events and the signal we are measuring must be necessarily weak, and the number of LECA families that can be traced back to specific ancestries is limited. Indeed the statistical significance using a Mann-Whitney U-test is often lost (>60%-70% of the times) when randomly reducing the data to sizes similar to the resulting sizes in their filtered dataset, which suggest that the mere effect of reducing the size, rather than the particular additional filtering used is having a major effect. This is why we made sure the signal was robust across different datasets, always using state of the art filtering approaches. Given the suggestion by Martin et. al. [1] that a recent phylogenetic analyses from them (which appeared after we had submitted the paper) represents a more careful dataset [3] we repeated our analyses using this dataset, which confirmed our results (650 eukaryotic clades, Archaeal vs Bacterial families, *P*=1.2e−41, two-tailed Mann-Whitney *U*-test and *α*-proteobacterial families’ *sl* significantly smaller within Bacterial, *P*=4.7e−2, permutation test, 10^6^ permutations). Again, this result lends further support to our findings.

Altogether, we show that the criticisms raised by Martin et. al. [1] do not comprise the main results and conclusions of our paper. Furthermore, we would like to stress that the new dataset and analyses brought about by this discussion lend additional support to our approach and conclusions.

## Methods

The analyses were based on the same alignment and tree data used in our original publication [2] and the data provided in [1]. All statistical tests were performed as in [2], using the same R and python code. For the new analyses, focusing on the various taxonomic levels within the LECA clades, the first encountered clade per level was accounted for the calculation of *rsl* and *ebl* values (analogously to the values of the LECA clades).

## References

[1] Martin, W. F. et al. Late mitochondrial origin is pure artefact. bioRxiv doi: http://dx.doi.org/10.1101/055368

[2] Pittis, A. A. & Gabaldón, T. Late acquisition of mitochondria by a host with chimaeric prokaryotic ancestry. Nature 531, 101–4 (2016).

[3] Ku, C. et al. Endosymbiotic origin and differential loss of eukaryotic genes. Nature 524, 427–432 (2015).

[4] Rasmussen, M. D. & Kellis, M. Accurate gene-tree reconstruction by learning gene- and species-specific substitution rates across multiple complete genomes. Genome Res. 17, 1932–42 (2007).

[5] Powell et. al. eggNOG v4.0: nested orthology inference across 3686 organisms. Nucleic Acids. Res. 42(Database issue):D231-9

